# Use of hiPSC-derived cardiomyocytes to rule out proarrhythmic effects of drugs: the case of hydroxychloroquine in COVID-19

**DOI:** 10.1101/2021.06.25.449913

**Authors:** Luca Sala, Vladislav Leonov, Manuela Mura, Alessandra Moretti, Lia Crotti, Massimiliano Gnecchi, Peter J Schwartz

## Abstract

In the early phases of the COVID-19 pandemic, drug repurposing was widely used to identify compounds that could improve the prognosis of symptomatic patients infected by SARS-CoV-2. Hydroxychloroquine (HCQ) was one of the first drugs used to treat COVID-19 patients due to its supposed capacity of inhibiting SARS-CoV-2 infection and replication *in vitro*.

While its efficacy is debated, HCQ has been associated with QT interval prolongation and potentially Torsades de Pointes, especially in patients predisposed to developing drug-induced Long QT Syndrome (LQTS) as silent carriers of variants associated with congenital LQTS. If confirmed, these effects represent a limitation to the at-home use of HCQ for COVID-19 infection as adequate ECG monitoring may be challenging.

We investigated the proarrhythmic profile of HCQ with Multi-Electrode Arrays after subchronic exposure of human induced pluripotent stem cell-derived cardiomyocytes (hiPSC-CMs) from two healthy donors, one asymptomatic and two symptomatic LQTS patients.

We demonstrate that: I) HCQ induced a concentration-dependent Field Potential Duration (FPD) prolongation *in vitro* and triggered arrhythmias that halted the beating at high concentration. II) hiPSC-CMs from healthy or asymptomatic carriers tolerated higher concentrations of HCQ and showed lower susceptibility to HCQ-induced electrical abnormalities regardless of baseline FPD values. These findings agree with the clinical safety records of HCQ and demonstrated that hiPSC-CMs potentially discriminates symptomatic vs asymptomatic mutation carriers through pharmacological interventions. Disease-specific cohorts of hiPSC-CMs may be a valid preliminary addition to quickly assess drug safety in vulnerable populations, offering rapid preclinical results with valuable translational relevance for precision medicine.

## Introduction

Drug repurposing is a key strategy aimed to identify new applications for compounds that have already been approved by regulatory authorities. The known safety profiles and toxicology of the repurposed compounds reduce the chances of failure and drug attrition, shorten the time frame for drug development and lower the costs associated with drug discovery and screening (Pushpakom et al., 2019). It had been a key process during the early phases of the COVID-19 pandemic to promote the identification of marketed compounds that could improve prognosis and therapy for symptomatic patients infected by the severe acute respiratory syndrome coronavirus 2 (SARS-CoV-2).

Hydroxychloroquine (HCQ), an antimalarial drug successfully used for the treatment of systemic lupus erythematosus and rheumatoid arthritis, was identified as a compound capable of inhibiting SARS-CoV-2 infection and replication *in vitro* (Liu et al., 2020; Touret et al., 2020; Yao et al., 2020); its off-label use was thus proposed for the treatment but also the prevention of COVID-19 in multiple clinical trials.

However, HCQ carries known pharmacological side-effects which include the block of the rapid delayed-rectifier potassium current (I_Kr_) and the inward delayed-rectifier potassium current (I_K1_) at therapeutic concentrations, with peak sodium current (I_Na_) and L-type calcium current (I_CaL_) blocked at higher concentrations (Wang et al., 2020).

At the clinical level, HCQ prolongs the QT interval (Saleh et al., 2020), with effects particularly exacerbated in the presence of factors such as plasma electrolyte imbalance (e.g. hypokalemia) and fever, both frequently present in patients with COVID-19, or secondary organ dysfunction. For these reasons, HCQ is classified by the CredibleMeds® database (Woosley et al.) as a drug with a known risk of causing Torsades de Pointes (TdP) and which has to be avoided in patients with the congenital Long QT Syndrome (cLQTS) (Schwartz and Ackerman, 2013). The proarrhythmic potential of HCQ seems further enhanced by its administration in combination with Azhithromycin, a macrolide antibiotic also proposed for the treatment of COVID-19 which can also cause TdPs (Mercuro et al., 2020; Fiolet et al., 2021).

Consequently, the administration of HCQ to a broad and heterogeneous population of patients, particularly at dosages higher than those already approved or while being proposed as at-home therapy regardless of any information on the patients’ genetic background (Procter et al., 2020), may create safety risks without adequate QT monitoring (Derwand et al., 2020); the susceptibility to QT-prolonging drugs is indeed exacerbated in case of silent cLQTS mutation carriers, i.e. patients carrying variant(s) in LQTS gene(s) who do not have the clinical hallmarks of LQTS despite having a compromised repolarization reserve (O’Hara and Rudy, 2012; Schwartz and Woosley, 2016). The potential proarrhythmic effect of HCQ has highlighted the complications of assessing the safety of pharmacological therapies, particularly on how to predict and quantify the risks of drugs used at doses not previously tested. A strategy to identify patients at risk of developing drug-induced QT prolongation and arrhythmias is still lacking, but there is a strong need for valid models which reproduce or predict the genotype-specific consequences of drugs on silent cLQTS mutation carriers; this would be particularly relevant for individuals with borderline QT intervals whose pathogenic variants often go undiagnosed in routine ECGs but which may significantly increase the arrhythmogenic susceptibility to QT prolonging drugs. This also applies for individuals carrying variants in protective/detrimental genetic modifiers (Itoh et al., 2016; Chai et al., 2018; Lee et al., 2021).

Despite the promising *in vitro* results, HCQ did not confirm its efficacy *in vivo* neither in preventing SARS-CoV-2 infection nor in treating the severe consequences of the infection, with findings confirmed by several independent studies (Graham et al., 2020; Axfors et al., 2021; Jorge, 2021; Macías et al., 2021; Rentsch et al., 2021; WHO Solidarity Trial Consortium et al., 2021). Conclusive evidence of its efficacy will come as several clinical trials are still recruiting (272 studies, 88 recruiting or enrolling by invitation; 24 Active, not recruiting; 9 suspended, 23 terminated; 62 completed; 28 withdrawn ClinicalTrials.gov, accessed on 14/04/2021). We believe that more information is required to understand the risks and boundaries of HCQ when administered in a potentially susceptible population of patients in which asymptomatic individuals carry a higher predisposition for TdP.

Here we used MultiElectrode Arrays (MEAs), one of the platforms of choice from the Comprehensive In vitro Proarrhythmia Assay (CiPA) consortium (Millard et al., 2018), to report the consequences of subchronic exposure to different concentrations of HCQ in a heterogeneous subset of three LQTS subjects and two unrelated healthy controls.

We aimed to assess whether: i) hiPSC-CMs can identify potential proarrhythmic effects by HCQ *in vitro*; ii) a genotype-specific response to HCQ can be reproduced with hiPSC-CMs and whether it correlates with clinical observations; and iii) hiPSC-CMs from homogeneous disease-specific cohorts could provide quick preclinical readouts before repurposing drugs to patients in large trials.

## Methods

### hiPSCs culture and differentiation to hiPSC-CMs

Three patients affected by LQTS were studied. One carries the KCNQ1-p.R594Q variant (Mura et al., 2018), associated with the most common form of LQTS (LQT1). One was affected by the Jervell and Lange-Nielsen syndrome (JLNS) carrying the KCNQ1-p.R594Q & KCNQ1-p.R190W variants in compound heteozigosity (Mura et al., 2018); JLNS is one of the most severe forms of LQTS, caused by the presence of homozygous or compound heterozygous variants on *KCNQ1* or on *KCNE1* associated to congenital deafness in addition to the typical cardiac features (Schwartz et al., 2006). One was affected by the CALM1-p.F142L variant (Crotti et al., 2013) associated with CALM-LQTS, which manifests in infants, is extremely severe, and responds poorly to therapy (Rocchetti et al., 2017; Crotti et al., 2019).

hiPSCs from healthy subjects: one was an unrelated *bona fide* healthy donor (WT (Lee et al., 2021) while hiPSCs from the second donor (WT2) were provided by the Coriell Institute for Medical Research (WTC-11 line, catalog No. GM25256).

hiPSCs were cultured on recombinant human vitronectin (rhVTN) in E8 Flex medium and differentiated on cell culture-grade Matrigel. hiPSCs were differentiated to hiPSC-CMs following a protocol based on the modulation of the Wnt-signalling pathway (Lian et al., 2012), purified through glucose starvation (> 90% CMs) and cryopreserved at days 17-20. Cryopreserved hiPSC-CMs were thawed before each experiment and maintained in culture for 7 days in RPMI medium supplemented with B27 Supplement (RPMI+B27). Data were obtained from 3 independent differentiations for each line. Details on reagents are provided in Table S1.

### MultiElectrode Array

Multiwell MEAs (MultiChannel Systems) were coated with 40 µg/mL bovine fibronectin and processed as previously described (Sala et al., 2017b). hiPSC-CMs were dissociated with TrypLE Select and replated on Multiwell MEAs as confluent monolayers. RPMI+B27 was refreshed every 48-72 hours. The recordings started after one week and were made at Baseline and at 2h, 24h and 48h after the treatment with HCQ (Figure 1).

**Figure 1.**
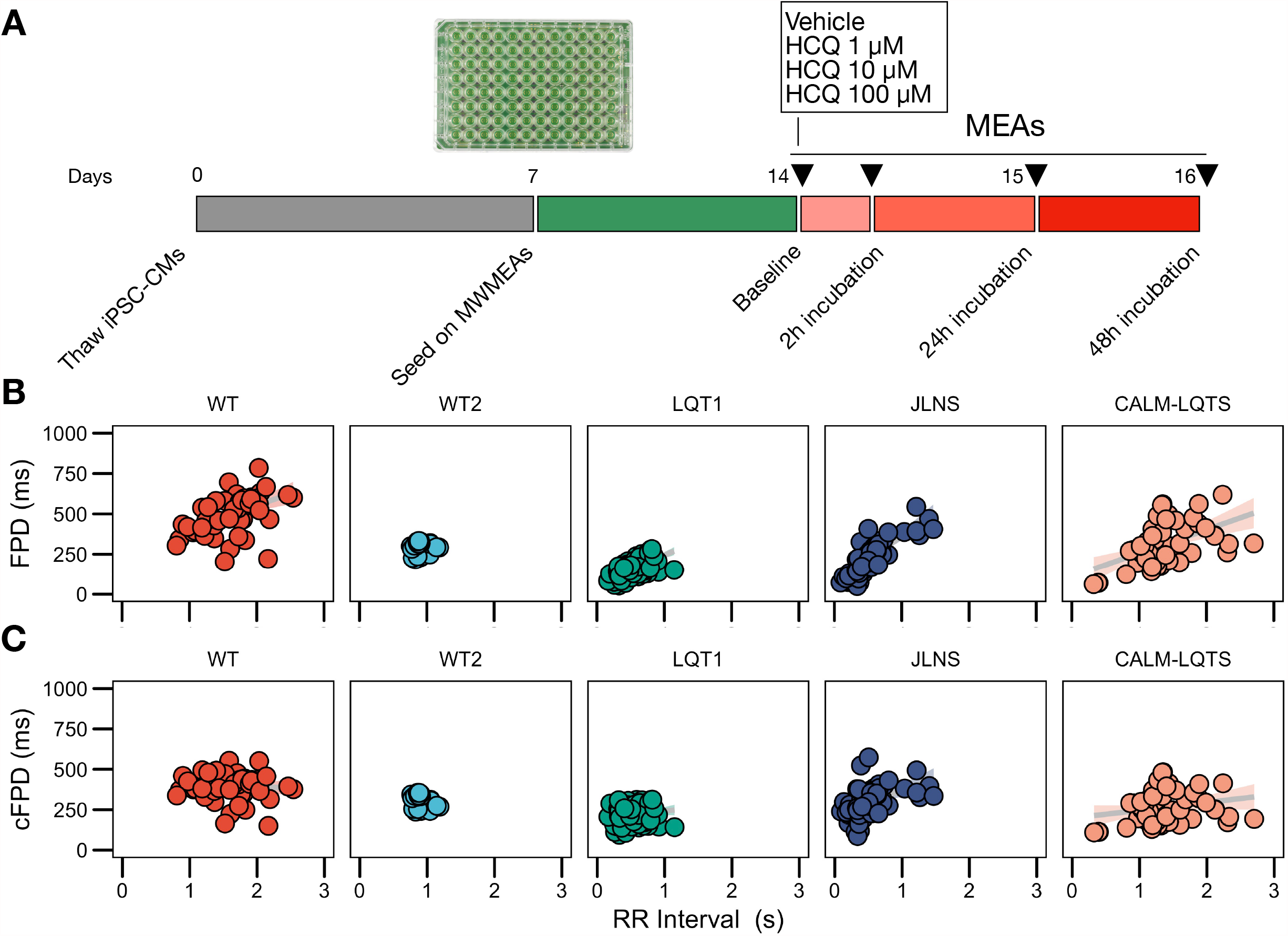
A) Study protocol. B) Relationships between FPD and RR for the iPSC lines used in this study measured with MWMEA at Baseline. Data were fitted with a linear model. N of MEA at Baseline: WT: 58; WT2: 48; LQT1: 106; JLNS: 62; CALM-LQTS: 56. C) Relationships between cFPD and RR for the iPSC lines used in this study measured with MWMEA at Baseline. Data were fitted with a linear model. N of MEA at Baseline: WT: 58; WT2: 48; LQT1: 106; JLNS: 62; CALM-LQTS: 56.

### Drug preparation and testing

HCQ was dissolved in water to obtain a stock solution of 50 mM. After having recorded Field Potentials (FP) values at Baseline, RPMI+B27 in Multiwell MEAs was refreshed with working concentrations of HCQ prepared in RPMI+B27 (1 µM, 10 µM, 100 µM). Vehicle was balanced accordingly and used as a control in each experiment and each Multiwell MEA plate.

### Field Potential Scoring System

A 6-point FP scoring system was implemented to evaluate the quality of FPs and to classify potential detrimental effects of the drug treatment on FP quality (Figure 3B).

Score 5: assigned to FPs with both markedly pronounced positive and negative upstrokes, a unambguous repolarization wave and a large FP amplitude; noise and signal oscillations at baseline must not be present and the RR interval must be regular.

Score 4: assigned to FPs with either a clear positive or negative depolarization peak, a visible repolarization wave and a large FP amplitude; noise and signal oscillations at baseline must not be present and the RR interval must be regular.

Score 3: assigned to FPs similar to the previous condition but with a smaller FP amplitude, a less markedly pronounced repolarization peak and a potentially ambiguous repolarization peak. Signal oscillations at baseline can be present and RR must be stable.

Score 2: assigned to FPs characterized by a small positive or negative depolarization, a small FP amplitude, a small and potentially ambiguous repolarization peak. Signal oscillations at baseline can be present and RR interval can be irregular.

Score 1: assigned to FPs characterized by a very irregular pattern(s), with small/absent depolarization and a small/absent repolarization peak; noise is present in terms of oscillations at baseline and electrical interference from neighboring electrodes, RR interval can be unstable.

Score 0: assigned to electrically inactive wells.

### Data Analysis and Statistics

The analysis of FPs was performed as previously described (Sala et al., 2017b) and automated through custom R scripts (R v.4.0.4) to accommodate a large number of data points. In total, data from 1320 individual MEA recordings were analyzed.

Comparisons of FPD, RR, cFPD were performed for each line with Dunnett’s test using Baseline as reference values. Statistical significance was defined as p<0.05 and is indicated in text and figures with an asterisk. Data are presented in the text as mean, mean ± standard error of the mean (sem) and scatter points where relevant.

To exclude any potential bias carried by the endogenous FPD of healthy hiPSC lines, we chose two healthy hiPSC lines characterized by different baseline FPD. Baseline FPD is a reliable parameter for comparisons only in case of isogenic lines or at least in case of lines sharing part of the genetic background (i.e. from relatives, as in case of LQT1 and JLNS lines in this paper) while it might often provide misleading results in case of cross-line comparisons (Sala et al., 2016, 2017a).

## Results

### Baseline data

Given the significant non-zero slope of the FPD-RR linear fit for the majority of the lines, data were corrected with the clinically-used Bazett’s QT correction formula, validated across all ages (Stramba-Badiale et al., 2018). Corrected FPDs (cFPD) nullified the positive correlation with RR where present (Figure 1B, C), normalizing the rate-dependency of FPD.

Signs of spontaneous arrhythmias and electrical instability were detected at baseline in MEAs from the JLNS and CALM-LQTS (Figure S2).

### Effect of HCQ on electrical activity and FP quality

We first investigated the presence of potential detrimental effects of the exposure to HCQ on the spontaneous activity of hiPSC-CMs monolayers.

We observed a dose-dependent effect of HCQ quantified as an increasing number of wells that ceased to beat with higher concentrations (Figure 2, Figure 3A). Here, a genotype-dependent effect was observed and it became particularly evident in two conditions: after the acute (2h) exposure to 10 µM HCQ, hiPSC-CMs from the two controls showed no effect on the spontaneous beating while ∼20% of the monolayers from the LQT1, ∼35% of the JLNS and ∼ 50% of the monolayers from the very severe CALM-LQTS temporarily halted the beating, which only partially recovered at longer timepoints (Figure 2). Similarly, 100 µM HCQ induced a larger termination of the spontaneous beating in monolayers from JLNS and CALM-LQTS, while asymptomatic LQT1 and the two WT seems more resistant to higher concentrations of HCQ (Figure 2 and 3A).

**Figure 2.**
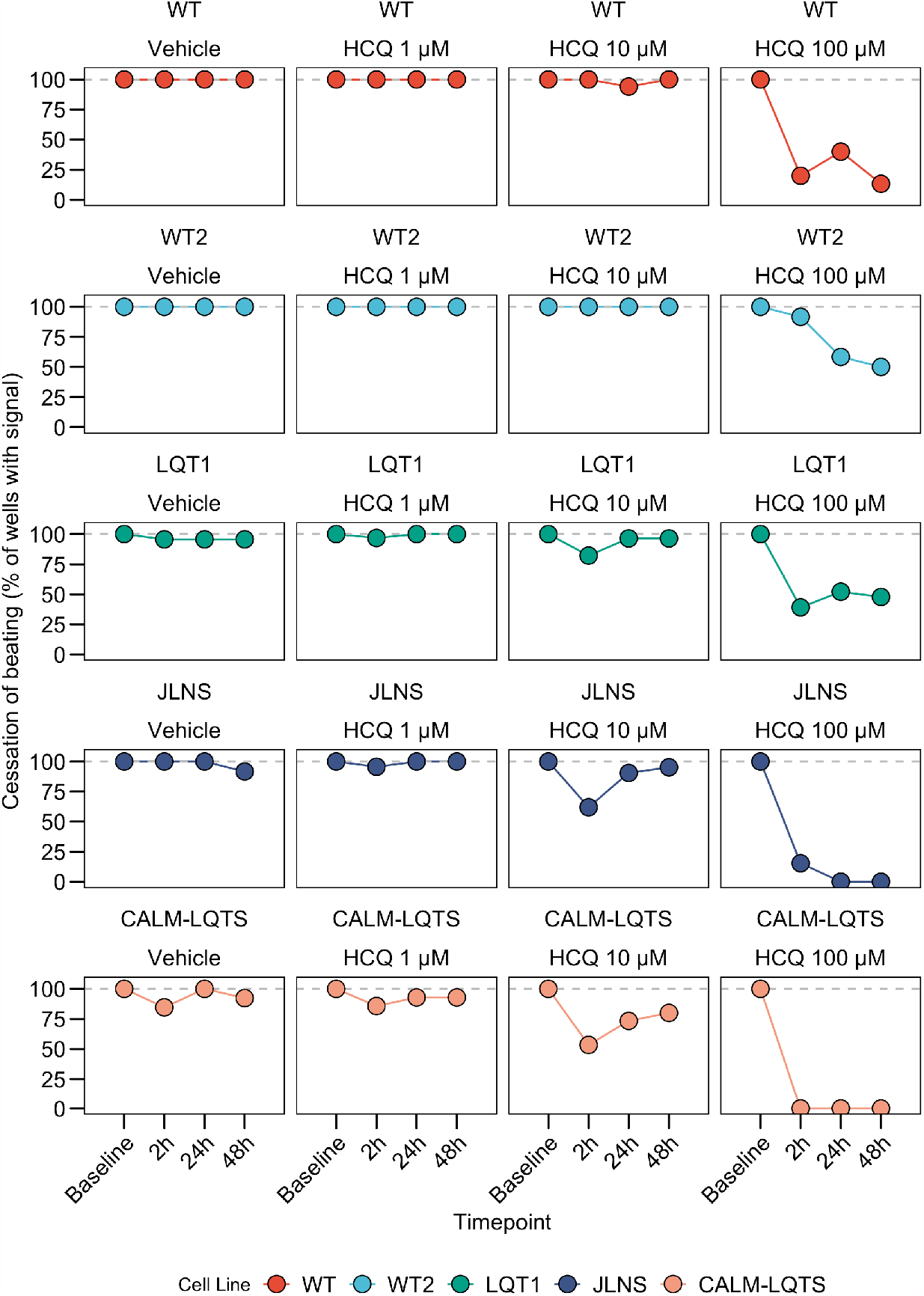
A) Cessation of beating, intended as the proportion of wells with a quantifiable signal at each timepoint.

**Figure 3.**
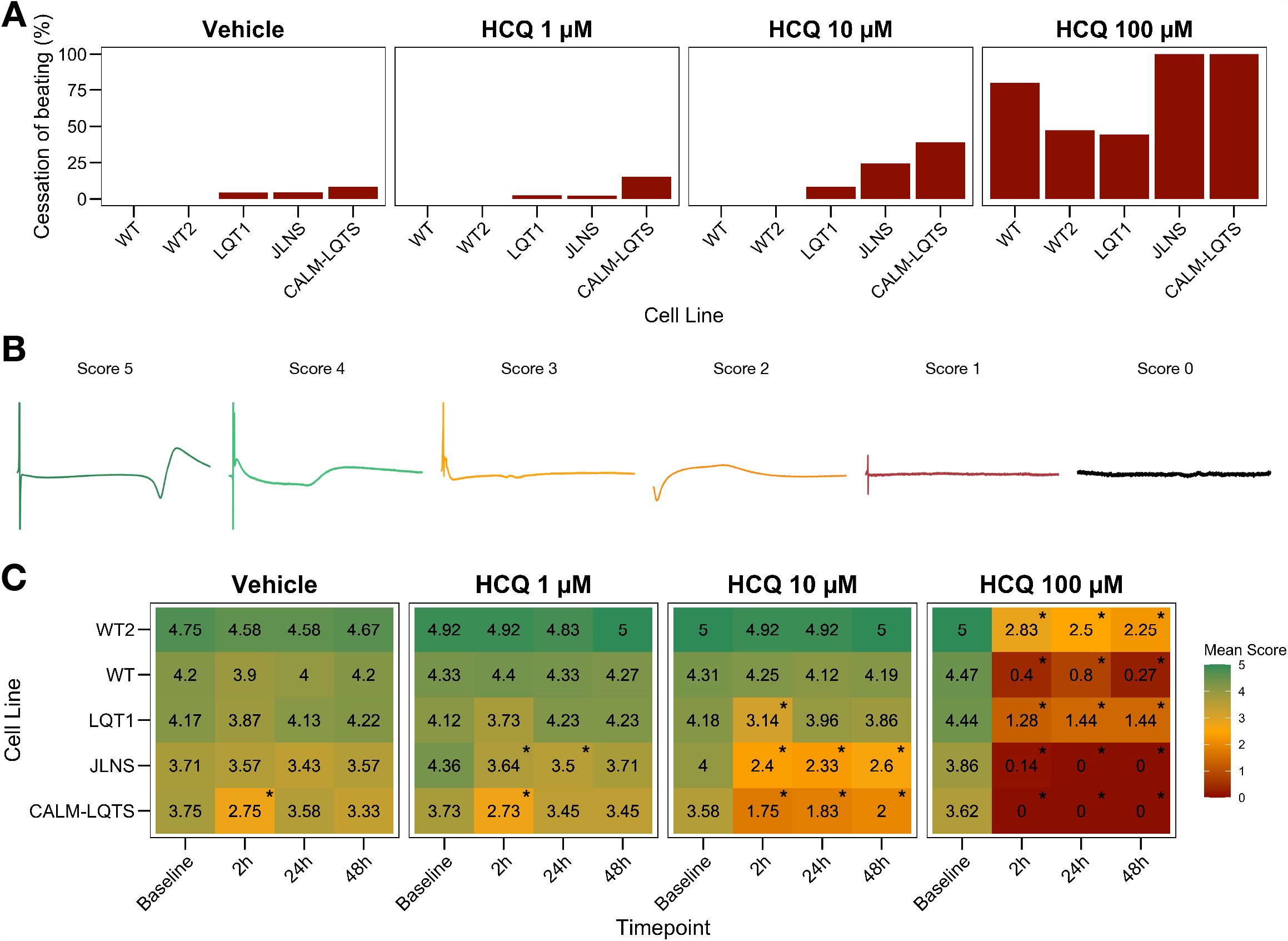
A) Percentage of MEAs that ceased to beat in at least one recording timepoint at different HCQ concentrations. B) Representative examples of FPs with the associated score that were used to quantify the effect of HCQ on FP quality. C) Mean FP scores at different HCQ concentrations and different timepoints in all the iPSC lines. The colour code indicates the mean FP score.

Next, we developed a 6-point scoring system to incorporate information regarding the FP quality degradation due to HCQ into the classification of the drug effect. This is particularly relevant to provide differential weights to MEAs in which the main temporal FP parameters (i.e. FPD, RR, cFPD, etc) could still be tracked and quantified but were spoiled by the drug treatment. We observed that HCQ had more severe effects on the FP quality of hiPSC-CMs from the two symptomatic subjects, with minor or modest effects in hiPSC-CMs from the asymptomatic LQT1 and from the healthy controls. These differences could be appreciated already at 1 µM and were particularly relevant at 10 µM. At 100 µM, only hiPSC-CMs from one of the two healthy donors maintained a higher proportion of good FPs despite their FP shapes being significantly modified.

### Effect of HCQ on FPD, RR, cFPD

After having assessed the detrimental effects of HCQ on FP quality, we investigated the effect of different concentrations of HCQ on MEA temporal parameters to verify whether HCQ could trigger a differential FPD prolongation based on the donor’s genotype. Higher susceptibility to media changes has been observed in the CALM-LQTS at all timepoints, even with vehicle, with spontaneous arrhythmic events often present (Figure S2). Treatment with vehicle induced changes (shortening) in the cFPD of WT at 2h, while it prolonged cFPD in CALM-LQTS at all the timepoints.

HCQ 1 µM induced prolongation in the cFPD of CALM-LQTS at all the timepoints and increased the number of MEAs without electrical activity.

HCQ 10 µM induced shortening in the cFPD of WT only at 2h, while it significantly increased cFPD in WT2, LQT1, JLNS and CALM-LQTS at all the time points and was paired with a cessation of beating only affecting LQT1, JLNS and CALM-LQTS.

HCQ 100 µM induced cFPD prolongation in WT, WT2, LQT1 while monolayers from JLNS and CALM-LQTS did not tolerate such high dosage and no electrical activity was observed at all timepoints.

## Discussion

Our findings provide novel evidence that hiPSC-CMs from subjects with a different propensity toward life-threatening arrhythmias, largely driven by their genetic background, may respond differently to drugs with the potential of blocking repolarizing currents. As drug screening moves progressively toward more sophisticated approaches, not unmindful of precision medicine, the incorporation of disease-specific cohorts of hiPSC-CMs should be considered as rational step in the assessment of drug repurposing strategies in vulnerable populations and possibly also in the safety screening of new drugs.

### Comparisons of concentrations between *in vitro and in vivo*

Although the concentrations and the modality of administration (acute vs cumulative) are different among *in vitro* experiments and many clinical trials, results for the WT subjects were similar to results from early timepoints published by other investigators with MEAs (TeBay et al., 2021) or in a more sophisticated organ-on-a-chip system (Charrez et al., 2021). Our study further extends these results including a susceptible population of patients affected by cLQTS that may be at higher risk of exhibiting diLQTS and developing TdP. The mean HCQ plasma concentration in subjects treated with HCQ has been measured as 50.3 ng/mL following a single 200 mg dose of HCQ (Plaquenil®) and a mean peak blood concentration of 129.6 ng/mL (Plaquenil label indication); based on these data, it is reasonable to estimate an *in vitro* concentration of 0.15 µM - 0.387 µM.

Inhibition of SARS-CoV-2 infection *in vitro* was achieved at concentrations larger than 4 µM (EC_50_ 4.06 µM or higher) regardless of the multiplicity of infection (MOI) used for SARS-CoV-2, (Liu et al., 2020), while the cytotoxic concentration (CC_50_) in ATCC-1586 cells was around 250 µM. Other studies identified a similar range of concentrations in Vero/VeroE6 cells, with an EC_50_ of ∼6 µM after 24h of treatment (Yao et al., 2020) or an IC_50_ of 2-4 µM at 48h or 72h respectively (Maisonnasse et al., 2020).

In the RECOVERY trial, patients were treated with 800 mg administered at zero and 6 hours, followed by 400 mg starting at 12 hours after the initial dose and subsequently every 12 hours for the following 9 days or until discharge (RECOVERY Collaborative Group et al., 2020). The simulated whole-blood concentration-time plots derived the theoretical whole-blood concentration of HCQ to span from 1 µM to 6 µM, with an upper safety bound for whole-blood of ∼10 µM (3 µM at plasma concentration) (White et al., 2020).

The concentrations used in this study appropriately covered clinically relevant concentrations as well as the *in vitro* effective dosages for SARS-CoV-2 inhibition, but also take into consideration a potential higher tolerance to drugs by hiPSC-CMs and the unknown effect of serum-free cell culture media on HCQ bioavailability.

### hiPSC-CMs can identify potential proarrhythmic effects of HCQ *in vitro*

Concentration-dependent effects were recorded for HCQ in all the lines, with the main modifications being changes in the repolarization peak as well as in the FP amplitude and quality. We could detect indications of proarrhythmic events in many of the MEAs treated with high HCQ concentrations, with the main events being abnormal shape of the repolarization wave, an irregular beat rate, a variable amplitude and presence of multiple depolarization peaks among consecutive beats. Some of these features were already visible at baseline for hiPSC-CMs from the two most severe lines used in this study (i.e. JLNS, CALM-LQTS).

### A genotype-specific response to HCQ can be reproduced with hiPSC-CMs and it correlates with the clinical severity of LQTS

A genotype-specific drug response pattern could also be identified and it aligned with the severity of the underlying genotypes. Our data indicate that hiPSC-CMs from the healthy donors or the asymptomatic LQT1 mutation carrier tolerated higher concentrations of HCQ and showed lower susceptibility to HCQ regardless of baseline *in vitro* FPD values.

Importantly and in agreement with previous observations (Sala et al., 2016, 2017a), the genotype-specific effects were present and predominant over the different baseline FPD parameters, further confirming that absolute FPD (or APD) values alone might provide misleading information when classifying drug effects in different cell lines, with the relative drug effects and a more comprehensive analysis of all FP parameters to take into account the underlying genotypes may provide more relevant results.

Healthy controls showed a higher tolerance for QTc prolongation and higher resistance to high HCQ concentrations. The asymptomatic LQT1 carrier (Female, Basal QTc = 458 ms) was identified in our centre only through a family screening of the sibling (JLNS, Female, Basal QTc = 578 ms, symptomatic) and might have been included in one of the COVID-19 clinical trials with HCQ because in most of them the exclusion criteria for QTc were a cutoff of 480 ms. This would have meant a significantly higher chance for a degradation of the electrical signal (Figure 3C) or for pathological FPD prolongation (Figure 4).

**Figure 4.**
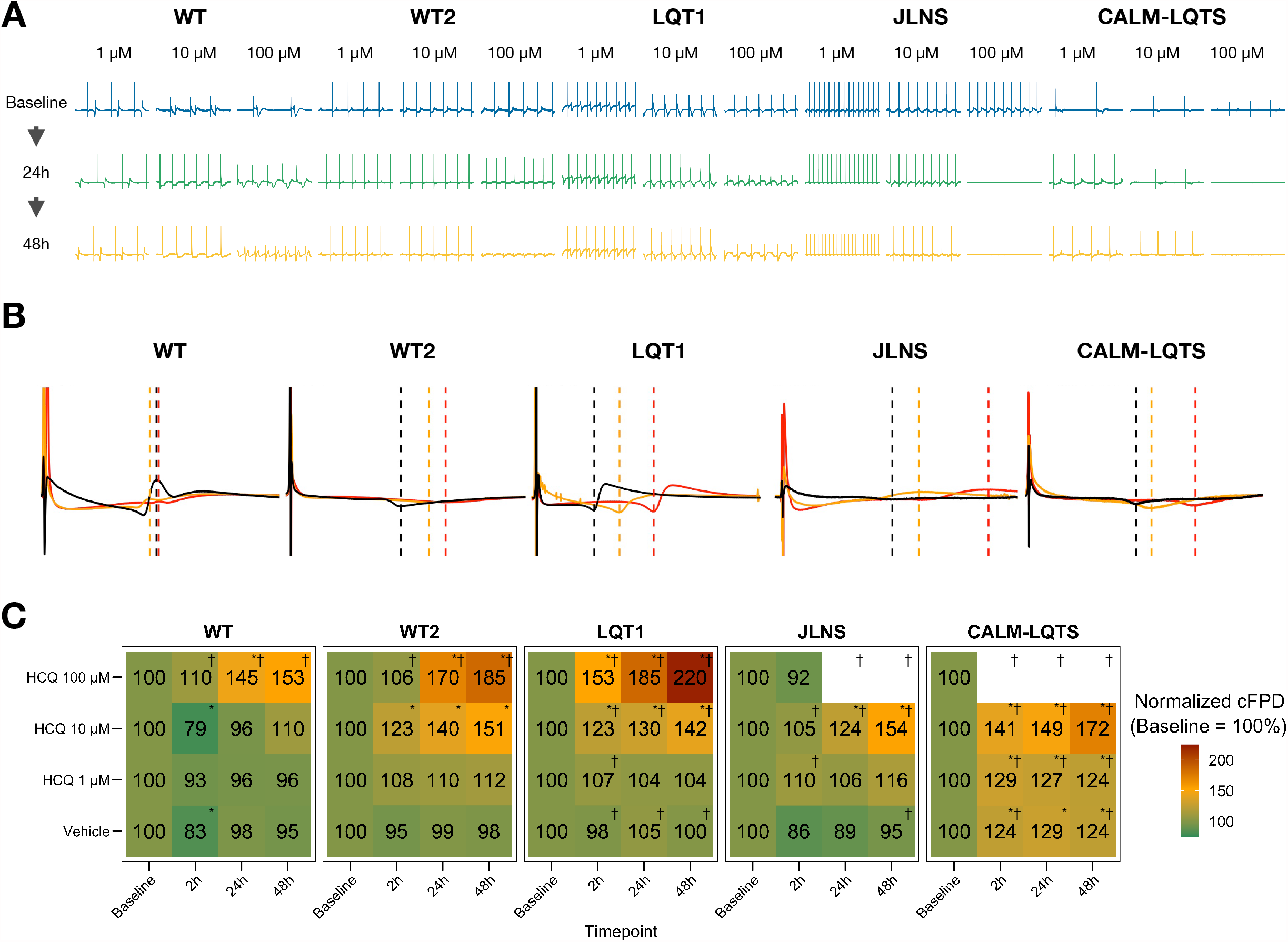
A) Representative examples of HCQ effect on MEA recordings. B) Representative mean FP profiles of MEAs treated with 10 µM HCQ at Baseline (black), after 24h (yellow) and 48h (red) exposure. C) Mean FPD change, expressed as % and normalized to the respective Baseline value. Colour code indicates the relative cFPD change compared to Baseline. * indicates p < 0.05 vs Baseline. † indicates that the treatment caused cessation of beating at a specific combination of Dosage and Timepoint.

Similarly to Chloroquine (Sánchez-Chapula et al., 2001), also HCQ blocks I_K1_, and this could explain the increased beating frequency observed particularly in hiPSC-CMs from WT despite HCQ should inhibit also the funny current (I_f_) (Capel et al., 2015); in our experimental settings, this effect did not clearly emerge in hiPSC-CMs from LQTS subjects; however, a drug-induced I_K1_ blockade may generate an additive effect on the underlying LQTS, especially on spontaneously beating cells, with the effect on the beating frequency potentially being masked by an additive and abnormal FPD prolongation due to an already compromized repolarization reserve.

### hiPSC-CMs offer rapid preclinical results with valuable translational relevance for precision medicine

Drug discovery and drug approval pipelines are already designed to prevent or minimize compounds that show a proarrhythmic potential due to a hERG blocking activity to be approved and commercialized (International Council for Harmonisation of Technical Requirements for Pharmaceuticals for Human Use, 2005). Even though important results have been achieved in the predictive potential of *in vitro* and *in silico* systems (Blinova et al., 2017; Li et al., 2017; Passini et al., 2017), comprehensive experimental models capable of recapitulating the complexity of human physiology are lacking and thus several compounds were withdrawn from the market after their approval (e.g. cisapride (Henney, 2000), astemizole (Gottlieb, 1999), terfenadine (Fung et al., 2001), etc), with cardiovascular toxicity still being a major cause for molecules to be discarded in the preclinical phases of drug development (Ferri et al., 2013). Platforms to guide clinical decision making and orient research efforts are required, particularly in critical situations when there is a compelling need for repurposed compounds or when therapies are being recommended for home use. In recent years, the use of hiPSC-CMs for safety pharmacology has progressively gained momentum and multiple validation strategies have been attempted to generate experimental models with enhanced predictive potential, and recently the CiPA approach or similar strategies have been used to evaluate the proarrhythmic risk of HCQ on hiPSC-CMs from commercial hiPSC lines or healthy individuals (Delaunois et al., 2021). Platforms based on hiPSCs can offer rapid and relevant readouts for safety pharmacology (Sala et al., 2017a) but the possibility of incorporating patient-specific drug responses in addition to those from healthy individuals can increase the predictive capacity of these platforms in conditions where prompt actions are required.

## Limitations of the study

Despite the remarkable progress in the strategies to mature hiPSC-CMs (Kamakura et al., 2013; Parikh et al., 2017; Feyen et al., 2020; Fukushima et al., 2020; Giacomelli et al., 2020), their electrophysiological phenotype still does not resemble that of adult cardiomyocytes and the relative ion channel distribution and drug responses might reflect this partial maturation; we anticipate that baseline phenotypes as well as raw dose-response curves might potentially differ in case other differentiation, purification or maturation strategies are used. Nevertheless, given the early expression of the main ion channel targeted by HCQ (i.e. Kv11.1), we expect the effects to be reproduced in other experimental settings; this work is in agreement with the longstanding clinical safety profile of HCQ in healthy individuals and also allowed solid drug-mediated discrimination of the underlying genotypes.

The intrinsic technical aspects of MEAs do not allow the comparison of absolute FP voltages as it would have been possible in single cells with patch clamp; this cost is repaid in terms of throughput. However, we cannot assume that the monolayers of hiPSC-CMs tested here, despite being constituted by purified populations, exhibited comparable resting membrane potentials and, thus, drug effects may be influenced by the inherent baseline electrophysiological properties of each line. This study provides information on the short-term (≤48h) arrhythmogenic potential of HCQ and this timeframe was sufficient to discriminate genotype-specific responses; this study does not provide information on cumulative effects caused by repetitive doses or long-term treatments.

## Conclusions

We have demonstrated that HCQ can induce cFPD prolongation, with effects becoming particularly important in susceptible subjects, at concentrations similar to those used in COVID-19 trials. More modest or negligible effects are instead evoked in hiPSC-CMs from healthy donors, particularly at concentrations close to those used to treat rheumatic disease; this was consistent with data from clinical studies (Abella et al., 2021).

Despite the highly controversial and debated (lack of) effectiveness for the prevention or treatment of COVID-19 (Axfors et al., 2021), the use of HCQ in a population of healthy individuals should not be stigmatized, as this compound has been successfully used for decades for the treatment of malaria, systemic lupus erythematosus and rheumatoid arthritis with more benefits than harm for patients (Wallace et al., 2012; Pareek et al., 2020), without significant increases in TdP risk at standard concentrations used in rheumatology (Taylor and White, 2004) and mainly only in association with co-prescribed QT-prolonging drugs and very high HCQ dosages (causal association) (Saint-Gerons and Tabarés-Seisdedos, 2021). Our report confirms this evidence.

Conversely, caution should be used for subjects who may be particularly susceptible to QT prolongation, and a particular focus should be put on silent or asymptomatic carriers of cLQTS variants and especially when genotype is additive to other detrimental pathophysiological factors or prescribed medications, as frequently occurs in patients with COVID-19 infection. Indeed, in a worldwide survey of COVID-19 associated arrhythmias, with almost 60% of the patients treated with hydroxychloroquine and 50% with azithromycin, those who developed cardiac arrhythmias already had borderline-prolonged QTc at the time of hospital admission and were more frequently affected by comorbidities and by severe pulmonary impairment (40% were mechanically ventilated) (Coromilas et al., 2021). On this issue, recommendations to further minimize risks of QT prolongation with Chloroquine or with HCQ in susceptible subjects have been recently drafted (Giudicessi et al., 2020; Offerhaus et al., 2020).

This work confirms that disease-specific hiPSC-CMs may provide additional information to drug screenings performed solely on hiPSC lines from healthy donors and may thus represent a viable option for preclinical drug repurposing screenings aiming for a higher translational readout. Larger homogeneous cohorts of hiPSC-CMs, with the ideal involvement of multiple facilities and including multiple disease-causing variants, genetic backgrounds and clinical manifestations, may represent a solid preliminary validation to assess the proarrhythmic effect of drugs, particularly in situations requiring rapid decisions. The combination of gene-editing and robust commercial iPSC lines from industry may further contribute to enhance and refine this aspect.

However, the most critical task remains the identification of who might be a susceptible subject and how a patient or a disease-cohort might respond to a new or repurposed therapy, and in this regard the contribution provided by subject-specific hiPSC-CMs combined with genome sequencing and *in silico* predictive approaches can provide insights with clinical implications for precision medicine (Gnecchi et al., 2021).

## Author Contributions

LS: study design, hiPSC-CMs culture, MEA, data analysis, wrote the manuscript.

VL: CALM1-p.F142L hiPSC-CMs culture, critical analysis of data and manuscript.

MM: LQT1-p.R594Q and JLNS-p.R594Q/p.R190W hiPSC line generation.

AM: CALM1-p.F142L hiPSC line generation.

LC: critical analysis of the manuscript.

MG: critical analysis of the manuscript.

PJS: critical analysis of the manuscript.

## Funding

This work was supported by a Marie Skłodowska-Curie Individual Fellowship (H2020-MSCA-IF-2017 No. 795209) to LS, by Fondazione CARIPLO, “Biomedical Research Conducted by Young Researchers” (grant No. 2019-1691) to LS, by a Leducq Foundation for Cardiovascular Research [18CVD05] ‘Towards Precision Medicine with Human iPSCs for Cardiac Channelopathies’ to PJS, MG, LC, LS, MM, by a PhD fellowship from the University of Verona to VL, by the DZHK and the German Research Foundation Transregio Research Units 152 and 267 to AM.

## Acknowledgments

The Authors would like to thank Dr. Michela Morano, PhD for assisting in hiPSCs culture and differentiation.

## Notes

### Competing Interest Statement

The authors have declared no competing interest.

