## Supplementary Information for "Use of hiPSC-derived cardiomyocytes to rule out proarrhythmic effects of drugs: the case of hydroxychloroquine in COVID-19"

**Table S1**
List of reagents used in the study.

| Reagent | Company | Product code |
|---|---|---|
| **Vitronectin (VTN-N) Recombinant Human Protein, Truncated** | Thermo Fisher Scientific [GIBCO] | A14700 |
| **Matrigel® Corning® hESC-Qualified Matrix, LDEV-free** | Corning | 354277 |
| **Fibronectin bovine plasma** | Merck [Sigma-Aldrich] | F1141 |
| **Essential 8™ Flex Medium Kit** | Thermo Fisher Scientific [GIBCO] | A2858501 |
| **RPMI 1640 w/ L-Glutamine** | Euroclone | ECB2000 |
| **RPMI 1640 Medium, no glucose** | Thermo Fisher Scientific [GIBCO] | 11879020 |
| **DMEM/F-12** | Thermo Fisher Scientific [GIBCO] | 11320033 |
| **B-27™ Supplement** | Thermo Fisher Scientific [GIBCO] | 17504044 |
| **B-27™ Supplement, minus insulin** | Thermo Fisher Scientific [GIBCO] | A1895601 |
| **DPBS, no calcium, no magnesium** | Thermo Fisher Scientific [GIBCO] | 14190094 |
| **EDTA (0.5 M), pH 8.0, RNase-free** | Thermo Fisher Scientific [Invitrogen] | AM9262 |
| **TrypLE™ Select Enzyme** | Thermo Fisher Scientific [GIBCO] | A1217701 |
| **RevitaCell™ Supplement** | Thermo Fisher Scientific [GIBCO] | A2644501 |
| **CryoStor® CS10** | StemCell Technologies | 7930 |
| **CHIR-99021 HCl** | Selleckchem | S2924 |
| **IWR-1** | Merck [Sigma-Aldrich] | I0161 |
| **24-well Plate with Gold Electrodes on FR4** | Multichannel Systems | 24W700/100F-288 |
| **96-well Plate with Gold Electrodes on FR4** | Multichannel Systems | 96W700/100F-288 |
| **Hydroxychloroquine Sulfate** | Selleckchem | S4430 |

**Table S2**
Baseline MEA parameters for all the lines used in the study.

| Cell Line | Mean FPD (ms) | FPD sem (ms) | Mean RR (ms) | RR sem (ms) | Mean cFPD (ms) | cFPD sem (ms) | N of MEAs |
|---|---|---|---|---|---|---|---|
| **WT** | 500.4 | 15 | 1639.8 | 50.5 | 393.9 | 9.9 | 58 |
| **WT2** | 286.1 | 5.1 | 890.5 | 12.3 | 304.1 | 5.7 | 48 |
| **LQT1** | 174.9 | 4.9 | 599.6 | 17.3 | 227.5 | 5.0 | 106 |
| **JLNS** | 225.5 | 13.6 | 546.4 | 37.1 | 303.9 | 11.7 | 62 |

| | | | | | | | |
|---|---|---|---|---|---|---|---|
| **CALM-LQTS** | 308.5 | 17.7 | 1359.9 | 62.6 | 263.7 | 13.1 | 56 |

### Supplementary Figures

**Figure S1**

A) Raw data for HCQ effect on FPD.
B) Raw data for HCQ effect on RR.
C) Raw data for HCQ effect on cFPD. In each plot, * indicates $p < 0.05$ vs Baseline.

**Figure S2**

A) Arrhythmogenic events detected in JLNS hiPSC-CMs.
B) Arrhythmogenic events detected in CALM-LQTS hiPSC-CMs.
