## Supplementary figures and images for "Use of hiPSC-derived cardiomyocytes to rule out proarrhythmic effects of drugs: the case of hydroxychloroquine in COVID-19"

### Figure S1

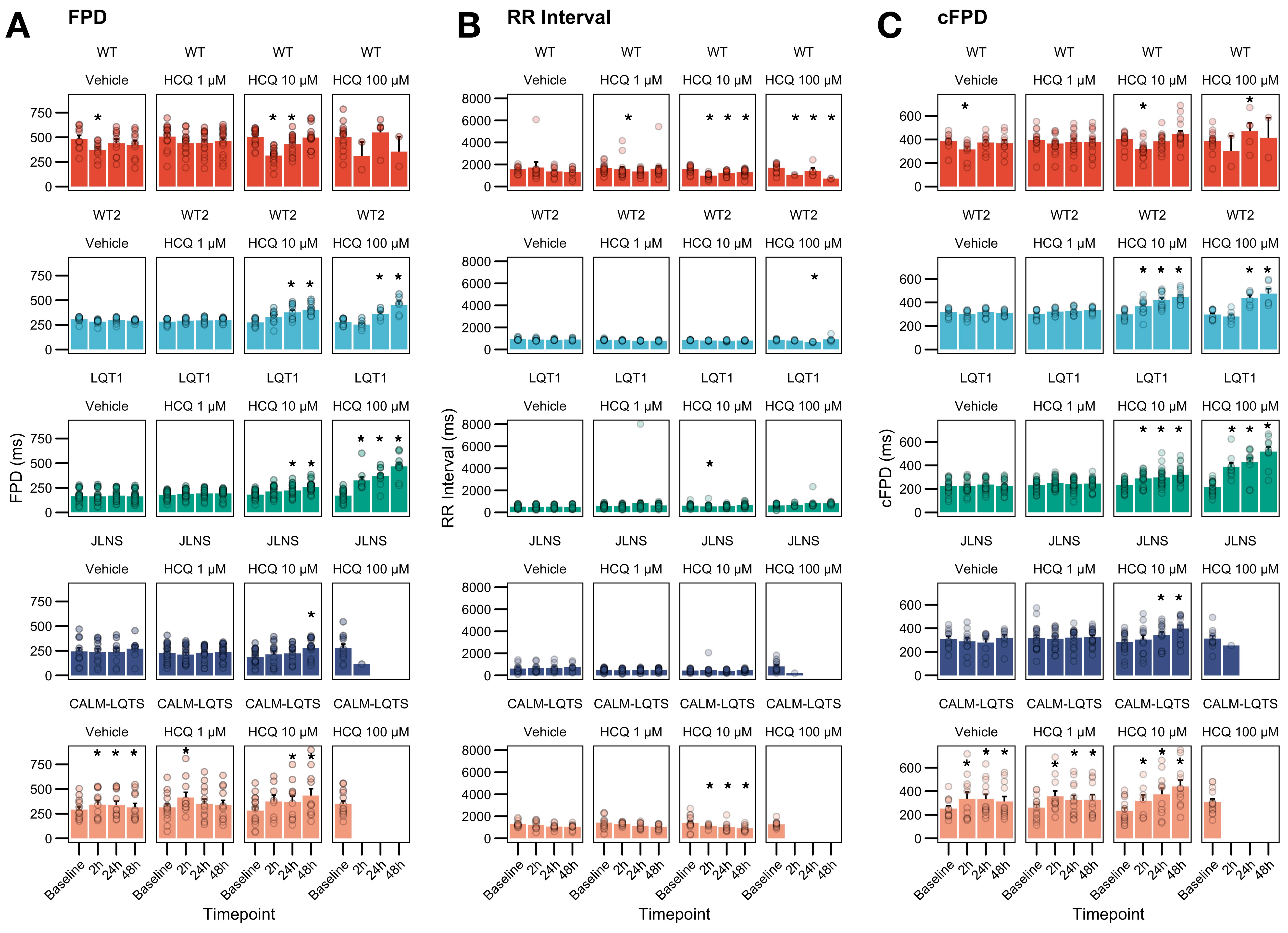

### Figure S2

#### A JLNS

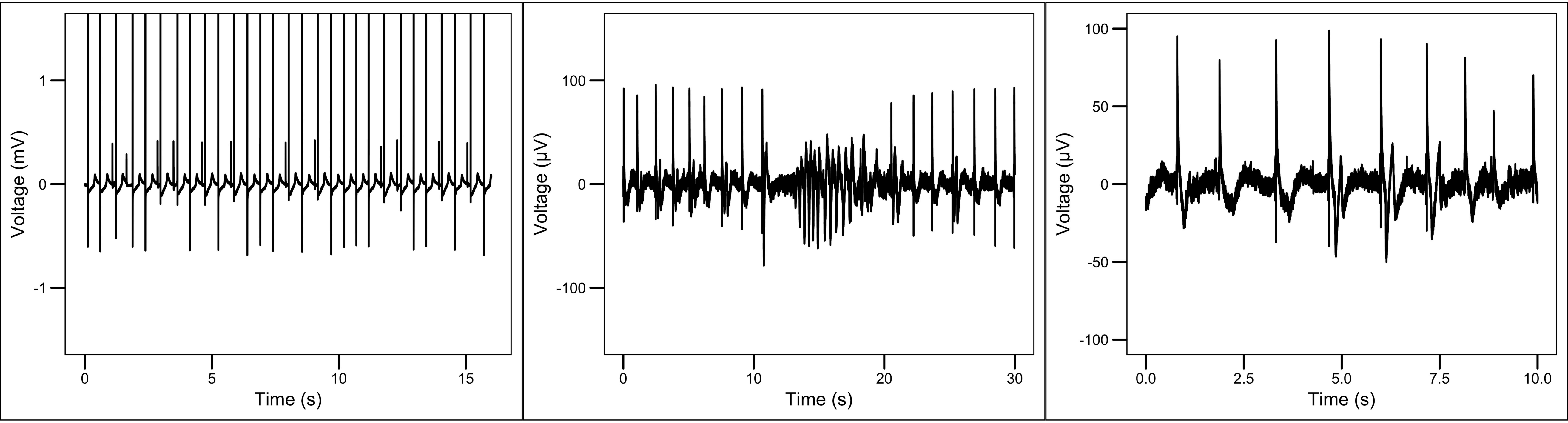


#### B CALM-LQTS

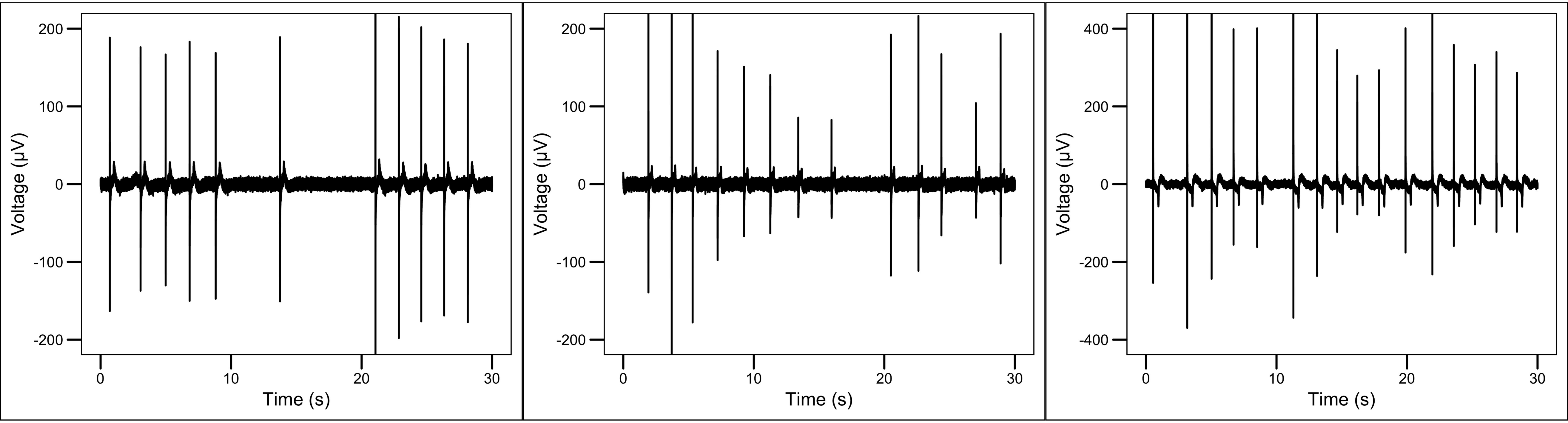
